# Characterization of emetic and diarrheal *Bacillus cereus* strains from a 2016 foodborne outbreak using whole-genome sequencing: addressing the microbiological, epidemiological, and bioinformatic challenges

**DOI:** 10.1101/399287

**Authors:** Laura M. Carroll, Martin Wiedmann, Manjari Mukherjee, David C. Nicholas, Lisa A. Mingle, Nellie B. Dumas, Jocelyn A. Cole, Jasna Kovac

**Author notes:** **Correspondence:** Jasna Kovac.

## Abstract

The *Bacillus cereus* group comprises multiple species capable of causing emetic or diarrheal foodborne illness. Despite being responsible for tens of thousands of illnesses each year in the U.S. alone, whole-genome sequencing (WGS) has not been routinely employed to characterize *B. cereus* group isolates from foodborne outbreaks. Here, we describe the first WGS-based characterization of isolates linked to an outbreak caused by members of the *B. cereus* group. In conjunction with a 2016 outbreak traced to a supplier of refried beans served by a fast food restaurant chain in upstate New York, a total of 33 *B. cereus* group strains were obtained from human cases (n =7) and food samples (n = 26). Emetic (n = 30) and diarrheal (n = 3) isolates were most closely related to *B. paranthracis* (clade III) and *B. cereus sensu stricto* (clade IV), respectively. WGS indicated that the 30 emetic isolates (24 and 6 from food and humans, respectively) were closely-related and formed a well-supported clade relative to publicly-available emetic clade III genomes with an identical sequence type (ST 26). When compared to publicly-available emetic clade III ST 26 *B. cereus* group genomes, the 30 emetic clade III isolates from this outbreak differed from each other by a mean of 8.3 to 11.9 core single nucleotide polymorphisms (SNPs), while differing from publicly-available genomes by a mean of 301.7 to 528.0 core SNPs, depending on the SNP calling methodology used. Using a WST-1 cell proliferation assay, the strains isolated from this outbreak had only mild detrimental effects on HeLa cell metabolic activity compared to reference diarrheal strain *B. cereus* ATCC 14579. Based on both WGS and epidemiological data, we hypothesize that the outbreak was a single source outbreak caused by emetic clade III *B. cereus* belonging to the *B. paranthracis* species. In addition to showcasing how WGS can be used to characterize *B. cereus* group strains linked to a foodborne outbreak, we also discuss potential microbiological and epidemiological challenges presented by *B. cereus* group outbreaks, and we offer recommendations for analyzing WGS data from the isolates associated with them.

## 1 Introduction

The *Bacillus cereus* (*B. cereus*) group, also known as *B. cereus sensu lato* (*s.l.*) is a complex of closely-related species that vary in their ability to cause disease in humans. Foodborne illness caused by members of the group primarily manifests itself in one of two forms: (i) emetic disease that is caused by cereulide, a heat-stable toxin produced by *B. cereus* within a food matrix prior to ingestion, or (ii) a diarrheal form of the disease, caused by enterotoxins produced in the small intestine of the host (Ehling-Schulz et al., 2004; Schoeni and Wong, 2005; Stenfors Arnesen et al., 2008). Here we refer to isolates that carry *ces* genes encoding the cereulide biosynthetic pathway as emetic isolates, and isolates that lack *ces* genes but carry either *hbl* or *cytK* genes that encode diarrheal enterotoxins as diarrheal isolates.

As foodborne pathogens, members of the *B. cereus* group are estimated to cause 63,400 foodborne disease cases per year in the U.S. (Scallan et al., 2011) and are confirmed or suspected to have been responsible for 235 outbreaks reported in the U.S. between 1998 and 2008 (Bennett et al., 2013). Due in part to its typically mild and self-limiting nature, foodborne illness caused by members of the *B. cereus* group is under-reported (Granum and Lund, 1997; Stenfors Arnesen et al., 2008), although severe infections resulting in patient death have been reported (Naranjo et al., 2011; Sanaei-Zadeh, 2012; Lotte et al., 2017). Furthermore, *B. cereus* group isolates that have been linked to human clinical cases of foodborne disease rarely undergo whole-genome sequencing (WGS), as is becoming the norm for other foodborne pathogens (Joensen et al., 2014; Ashton et al., 2015; Moura et al., 2017).

Here, we describe a foodborne outbreak caused by members of the *B. cereus* group in which WGS was implemented to characterize isolates from human clinical cases and food. To our knowledge, this is the first description of a *B. cereus* outbreak in which WGS was employed to characterize isolates. By testing various combinations of variant calling methodologies, we showcase how different bioinformatics pipelines can yield vastly different results when pairwise SNP differences are the desired metric for determining whether an isolate is part of an outbreak or not. In addition to discussing the bioinformatic challenges, we examine potential microbiological and epidemiological obstacles that can hinder characterization of *B. cereus* group isolates from suspected foodborne outbreaks, and we offer recommendations to guide the characterization of future *B. cereus* group outbreaks using WGS.

## 2 Materials and Methods

### 2.1 Collection of epidemiological data

Epidemiological investigations were coordinated by the New York State Department of Health (NYSDOH), and the outbreak was reported to the U.S. Centers for Disease Control and Prevention (CDC). Investigation methods included (i) a cohort study, (ii) food preparation review, (iii) an investigation at a factory/production/treatment plant, (iv) food product traceback, and (v) environment/food/water sample testing.

### 2.2 Isolation and initial characterization of *B. cereus* strains

Stool specimens were plated directly onto mannitol-egg yolk-polymyxin (MYP) agar and incubated aerobically at 37°C for 24 h. Food samples were diluted 1:10 in 1 X PBS, pH 7.4 in a filter bag for homogenizer blenders and homogenized for 2 min. One hundred μl of each homogenized sample were plated onto MYP agar and incubated aerobically at 37°C for 24 h. The MYP agar plates for both the stool specimens and food samples were observed after the 24-hour incubation period. Individual *B. cereus*-like colonies (i.e., pink colored and lecithinase positive) were subcultured to trypticase soy agar (TSA) plates supplemented with 5% sheep blood and incubated aerobically at 37°C for 18-24 h. These isolates were identified as *B. cereus* using the following conventional microbiological techniques: Gram stain, colony morphology, hemolysis, motility, and spore stain. To differentiate between *B. cereus* and *B. thuringiensis*, isolates were cultured for 48 h at 37°C on sporulation agar slants. Smears were prepared, and slides were heat fixed and then stained using malachite green and counter stained with carbol fuchsin (Tallent et al., 2012). Slides were then observed for the presence or absence of parasporal crystals.

### 2.3 *rpoB* allelic typing

The 33 outbreak isolates were streaked onto brain heart infusion (BHI) agar from their respective cryo stocks stored at −80 °C and incubated overnight at 37 °C. Single isolated colonies were inoculated in 5 ml BHI broth and incubated overnight at 32 °C and used for genomic DNA extraction using Qiagen DNeasy blood and tissue kits (Qiagen). Extracted DNA was used as a template in a PCR reaction using primers targeting a 750 bp sequence of the *rpoB* gene (RzrpoBF: AARYTIGGMCCTGAAGAAAT and RZrpoBR: TGIARTTRTCATCAACCATGTG) (Ivy et al., 2012). PCR was carried out in 25 μl reactions using GoTaq Green Master Mix (Promega Corporation) under the following thermal cycling conditions: 3 min at 94°C, followed by 40 cycles of 30 s at 94°C, 30 s at 55-45°C (in the first 20 cycles the temperature was reduced for 0.5°C per cycle and then kept at 45°C in the following 20 cycles), followed by 1 min at 72°C, and a final hold at 4°C. The resulting PCR product was used for genotyping and preliminary species identification (Ivy et al., 2012).

### 2.4 Bacterial growth conditions and collection of bacterial supernatants

The 33 outbreak isolates, as well as *B. cereus s.s.* type strain ATCC 14579 and *B. cereus* emetic reference strain DSM 4312 (Food Microbe Tracker ID FSL M8-0547; Vangay et al., 2013) were streaked onto BHI agar from their respective cryo stocks stored at −80 °C. Single isolated colonies were inoculated in 5 ml BHI broth and incubated at 37 °C without shaking. For immunoassays and cytotoxicity assays (see sections 2.5 and 2.6), overnight cultures (grown for 18 h at 37 °C) were used for inoculation of fresh BHI broth, and the cultures were grown to early stationary phase (OD_600_ of approximately 1.5, which equals approximately 10^8^ CFU/ml). The growth was quenched by placing them on ice. The cultures were spun down at 16,000 g for 2 min, and the supernatants were collected, aliquoted in duplicate, and stored at −80°C until further use.

### 2.5 Hemolysin BL and nonhemolytic enterotoxin detection

Diarrheal strains grown as described above were used for qualitative detection of hemolysin BL (Hbl) and nonhemolytic enterotoxins (Nhe) with the Duopath Cereus Enterotoxins immunoassay (Merck). Only select representatives of emetic outbreak strains were tested (i.e., FSL R9-6381, FSL R9-6382, FSL R9-6384, FSL R9-6389, FSL R9-6395, and FSL R9-6399), as they did not carry genes encoding Hbl and were therefore not expected to produce Hbl. Briefly, the temperature of cultures and immunoassays was adjusted to room temperature. 150 µl of each isolate culture were added to the immunoassay port, following the manufacturer’s instructions. The results were read as positive if a red line was visible after a 30-min incubation at room temperature. Tests were considered valid only when controls lines were visible.

### 2.6 WST-1 metabolic activity assay

HeLa cells were seeded in 96-well plates at a seeding density of 8 × 10^4^ cells/cm^2^ (Fisichella et al., 2009) in Eagle’s minimum essential medium (EMEM) supplemented with 10% fetal bovine serum (FBS) and allowed to grow for 18-24 h at 37°C, 5% CO_2_. The medium in each well was replaced with 100 µl of fresh medium containing 5% v/v of bacterial supernatants (prepared as described above) that were thawed, pre-warmed to 37°C, and mixed. The medium containing supernatants was added to the cells using a multichannel pipettor to minimize the variability in the duration of cell exposure to the toxin amongst wells of a 96-well plate. Medium containing 5% BHI was used as a negative control and medium containing 5% of 1% Triton X-100 prepared in BHI (final concentration in the well of 0.05%) was used as a positive control, with the latter expected to reduce the viability of HeLa cells. After 15 min of intoxication at 37 °C, 5% CO_2_ (Miller et al., 2018), 10 µl of WST-1 dye solution (Roche) was added to each well of the plate, and the plate was incubated for 25 min at 37 °C, 5% CO_2_, resulting in a total of 40 min exposure of cells to the supernatants. After 30 s of orbital shaking at 600 rpm, the absorbances were read by a microplate reader (Thermo Scientific Multiskan GO, Thermo Fisher Scientific) in precision mode at 450 nm and 690 nm, the latter being subtracted from the former to account for the background signal (i.e., corrected absorbances) (Fisichella et al., 2009). Each test, including 0.05% Triton X-100, was conducted with six technical replicates and on two different HeLa passages using supernatants from single biological replicates, resulting in a total of 12 technical replicates per isolate. The viability of cells was determined by calculating a ratio of corrected absorbances to that of BHI, converting to percentages, and calculating the mean of technical replicates for each isolate. The results were compared to the results for cells treated with (i) 0.05% Triton X-100, (ii) *B. cereus s.s.* type strain ATCC 14579 supernatant (i.e., reference for diarrheal strains), and (iii) *B. cereus* group strain DSM 4312 supernatant (i.e., reference for emetic strains).

### 2.7 Statistical analysis of cytotoxicity data

A Welch’s test and the Games-Howell post-hoc test (appropriate for data with non-homogeneous variances) were performed using results of all 12 technical replicates of each outbreak-associated isolate, as well as on *B. cereus s.s.* type strain ATCC 14579, emetic reference stain *B. cereus* DSM 4312, and 0.05% Triton X-100. For the Games-Howell test, a Bonferroni correction was applied to correct for multiple comparisons. Statistical analyses were carried out in R version 3.4.3 (R Core Team, 2018).

### 2.8 Whole-genome sequencing

Genomic DNA was extracted from overnight cultures (∼18 h) grown in BHI at 32°C using Qiagen DNeasy blood and tissue kits (Qiagen) or the Omega E.Z.N.A. Bacterial DNA kit (Omega) following the manufacturers’ instructions. For the E.Z.N.A. Bacterial DNA kit, the additional steps recommended for difficult to lyse bacteria were taken to obtain sufficient DNA yield. Briefly, one ml of an overnight culture was additionally treated with glass beads provided in the E.Z.N.A. kit. DNA was quantified using Qubit 3 and used for Nextera XT library preparation (Illumina). Pooled libraries were sequenced in two Illumina sequencing runs with 2 × 250 bp reads at the Penn State Genomics Core Facility and at the Cornell Animal Health Diagnostic Center.

### 2.9 Initial data processing and genome assembly

Illumina adapters and low-quality bases were trimmed using Trimmomatic version 0.36 (Bolger et al., 2014) for Nextera paired-end reads, and FastQC version 0.11.5 www.bioinformatics.babraham.ac.uk/projects/fastqc/) was used to confirm that read quality was adequate. Genomes listed in Supplementary Table S1 were assembled *de novo* using SPAdes version 3.11.0 (Bankevich et al., 2012), and average per-base coverage was calculated using BWA MEM version 0.7.13 (Li and Durbin, 2010) and Samtools version 1.6 (Li et al., 2009).

### 2.10 *In silico* typing and virulence gene detection

BTyper version 2.2.0 (Carroll et al., 2017) was used to perform *in silico* virulence gene detection, multi-locus sequence typing (MLST), *panC* clade assignment, and *rpoB* allelic typing, as well as to extract the gene sequences for all detected loci. For virulence gene detection, the default settings were used (i.e., 50% amino acid sequence identity, 70% query coverage), as these cut-offs have been shown to correlate with PCR-based detection of virulence genes in *B. cereus* group isolates (Kovac et al., 2016; Carroll et al., 2017). BMiner version 2.0.2 (Carroll et al., 2017) was used to aggregate the output files from BTyper and create a virulence gene presence/absence matrix.

### 2.11 Construction of *k*-mer based phylogeny using outbreak strains and genomes of 18 *B. cereus* group species

kSNP version 3.1 (Gardner and Hall, 2013; Gardner et al., 2015) was used to produce a set of core SNPs among the 33 outbreak genomes, plus genomes from each of the 18 *B. cereus* group species listed in Supplementary Table S2, using the optimal *k*-mer size as determined by Kchooser (*k* = 21). The resulting core SNPs were used in conjunction with RAxML version 8.2.11 (Stamatakis, 2014) to construct a maximum likelihood (ML) phylogeny using the GTRCAT model with a Lewis ascertainment bias correction (Lewis, 2001) and 500 bootstrap replicates. The resulting phylogenetic tree was formatted using the phylobase (R Hackathon et al., 2017), ggtree (Guangchuang et al., 2017), phytools (Revell, 2012), and ape (Paradis et al., 2004) packages in R version 3.4.3.

### 2.12 Variant calling and phylogeny construction using outbreak isolates

Combinations of five reference-based variant calling pipelines (Table 1) and reference genomes (Table 2), as well as one reference-free SNP calling pipeline (Table 1), were used to separately identify core SNPs among (i) all 33 outbreak-related isolates (30 emetic clade III isolates and 3 clade IV isolates) and (ii) the subset of 30 emetic clade III isolates.

**Table 1.** Variant calling pipelines tested in this study.

| Pipeline<sup>a</sup> | Approach | Reference-based | Input data (file format)<sup>b</sup> | Read mapper | Variant caller | Reference(s) and in-depth pipeline descriptions |
| --- | --- | --- | --- | --- | --- | --- |
| **CFSAN** | Read mapping | Yes | PE reads (fastq) | Bowtie2 | Varscan | http://snp-pipeline.readthedocs.io/en/latest/ |
| **Freebayes** | Read mapping | Yes | PE reads (fastq) | BWA MEM | Freebayes | https://github.com/lmc297/SNPBac |
| **kSNP3** | *k*-mer based | No | Contigs (fasta) | Not applicable | kSNP3 | https://sourceforge.net/projects/ksnp/files/ |
| **LYVE-SET** | Read mapping | Yes | PE reads (fastq) | SMALT | Varscan | https://github.com/lskatz/lyve-SET |
| **Parsnp** | Core genome alignment | Yes | Contigs (fasta) | Not applicable | Parsnp | http://harvest.readthedocs.io/en/latest/content/parsnp.html |
| **Samtools** | Read mapping | Yes | PE reads (fastq) | BWA MEM | Samtools/Bcftools | https://github.com/lmc297/SNPBac |

<sup>a</sup>CFSAN, U.S. Food and Drug Administration (FDA) Center for Food Safety and Applied Nutrition SNP pipeline; LYVE-SET, U.S. Centers for Disease Control and Prevention (CDC) *Listeria*, *Yersinia*, *Vibrio*, and *Enterobacteriaceae* SNP Extraction Tool
<sup>b</sup>PE reads, Illumina paired-end reads

**Table 2.** Reference genomes used for reference-based variant calling in this study.

| Reference Genome | Clade<sup>a</sup> | Data set(s)<sup>b</sup> | ANI Range<sup>c</sup> | NCBI<br>Accession | Assembly<br>Level | Rationale for Selection |
| --- | --- | --- | --- | --- | --- | --- |
| *B. cereus* strain ATCC 14579 chromosome | IV | All 33 isolates from two clades (clades III and IV) | 98.8-98.9 (clade IV)<br>91.8-92.3 (clade III) | NC_004722.1 | Complete Genome | *B. cereus* s.s. type strain; RefSeq reference genome; member of *panC* clade IV, the same clade as the three non-emetic outbreak-associated isolates sequenced in this study |
| *B. cereus* strain AH187 chromosome | III | All 33 isolates from two clades (clades III and IV); 30 emetic clade III isolates | 92.0-92.2 (clade IV)<br>99.8-99.9 (clade III) | NC_011658.1 | Complete Genome | Human clinical isolate associated with an emetic outbreak in 1972 (cooked rice, United Kingdom); identical virulotype, MLST sequence type, *rpoB* allelic type, and *panC* clade as 30 emetic outbreak isolates sequenced in this study |
| *B. cytotoxicus* strain NVH 391-98 chromosome | VII | All 33 isolates from two clades (clades III and IV) | 82.6-82.7 (clade IV)<br>82.5-82.9 (clade III) | NC_009674.1 | Complete Genome | Type strain of *B. cytotoxicus*, the most distant member of the *B. cereus* group as currently defined; shares a common ancestor with all isolates sequenced in this study |
| FOOD_10_19_16_RSNT1_2H_R9-6393 | III | 30 emetic clade III isolates | 92.0-92.2 (clade IV)<br>100<sup>d</sup>-100 (clade III) | SRR6825038 | Contigs | Emetic isolate from the outbreak reported here; assembly had high per-base coverage, as well as the fewest number of contigs of all genome assemblies from isolates in this outbreak |

<sup>a</sup>Clade determined via *panC* clade assignment function in BTyper version 2.2.0
<sup>b</sup>Data set(s) in this study for which a given genome was used as a reference genome for reference-based SNP calling
<sup>c</sup>Minimum and maximum average nucleotide identity (ANI) values of reference strain relative to clade IV and clade III genomes sequenced in this outbreak (n = 3 and 30, respectively) calculated using FastANI
<sup>d</sup>Minimum ANI value was less than 100 prior to rounding

For the Samtools and Freebayes pipelines (Table 1), trimmed Illumina paired-end reads from the queried isolates were mapped to the appropriate reference genome using BWA mem version 0.7.13 (Li, 2013) and either Samtools/Bcftools version 1.6 (Li et al., 2009) or Freebayes version 1.1.0 (Garrison and Marth, 2012), respectively, were used to call variants. vcftools version 0.1.14 (Danecek et al., 2011) was used to remove indels and SNPs with a mapping quality score < 20, as well as to construct consensus sequences. For both variant calling pipelines, Gubbins version 2.2.0 (Croucher et al., 2015) was used to filter out recombination events from the consensus sequences. Both of these pipelines are publicly-available and can be reproduced in their entirety (SNPBac version 1.0.0; https://github.com/lmc297/SNPBac).

For the CFSAN (Davis et al., 2015) and LYVE-SET (Katz et al., 2017) pipelines (versions 1.0.1 and 1.1.4g, respectively; Table 1), trimmed Illumina paired-end reads were used as input, and all default pipeline steps were run as outlined in the manuals. For the Parsnp pipeline (Treangen et al., 2014) (Table 1), assembled genomes of the outbreak isolates were used as input, and Parsnp’s implementation of PhiPack (Bruen et al., 2006) was used to filter out recombination events. For kSNP3 (Table 1), assembled genomes of the outbreak isolates were used as input, and Kchooser was used to determine the optimum *k*-mer size for the full 33-isolate data set and the 30 emetic clade III isolate set (*k* = 21 and 23, respectively).

For all variant calling and filtering pipelines, RAxML version 8.2.10 was used to construct ML phylogenies using the resulting core SNPs under the GTRGAMMA model with a Lewis ascertainment bias correction and 1,000 bootstrap replicates. Phylogenetic trees were annotated using FigTree version 1.4.3 (http://tree.bio.ed.ac.uk/software/figtree/).

### 2.13 Variant calling and statistical comparison of emetic outbreak isolates to publicly-available genomes

To compare emetic clade III isolates from this outbreak to other emetic clade III isolates, BTyper version 2.2.1 was used to query all 2,156 *B. cereus* group genome assemblies available in NCBI’s RefSeq database (Pruitt et al., 2007) and identify all genome assemblies that (i) belonged to clade III based on *panC* sequence, (ii) belonged to ST 26 using *in silico* MLST, and (iii) were found to possess the *ces* operon in its entirety (*cesABCD*) at the default coverage and identity thresholds. This search produced 25 genome assemblies in addition to the 30 emetic clade III genomes sequenced here. Only three of the 25 RefSeq genome assemblies had Sequence Read Archive (SRA) data linked to their BioSample accession numbers, making short read data readily available only for these three isolates. Consequently, only Parsnp version 1.2 and kSNP version 3.1 were used to identify SNPs in all 55 clade III emetic genomes (25 from NCBI RefSeq and 30 sequenced here), as these approaches can be used with assembled genomes and do not require short reads as input. For Parsnp, the chromosome of *B. cereus* str. AH187 was used as a reference genome. For kSNP3, Kchooser was used to select the optimal *k*-mer size (*k* = 21), and the chromosome of *B. cereus* str. AH187 was included for *k*-mer based SNP calling.

RAxML version 8.2.10 was used to construct ML phylogenies using the resulting core SNPs for each of the Parsnp and kSNP3 pipelines under the GTRCAT model with a Lewis ascertainment bias correction and 1,000 bootstrap replicates. Pairwise core SNP differences between all 55 isolates were obtained using the dist.gene function in R’s ape package. The permutest and betadisper functions in R’s vegan package (Oksanen et al., 2018) were used to conduct an ANOVA-like permutation test to test if publicly-available genomes were more variable than isolates from this outbreak based on pairwise core SNP differences and 5 independent trials using 100,000 permutations each. Analysis of similarity (ANOSIM; Clarke, 1993) using the anosim function in the vegan package in R was used to determine if the average of the ranks of within-group distances was greater than or equal to the average of the ranks of between-group distances (Anderson and Walsh, 2013), where groups were defined as (i) the 30 emetic isolates from this outbreak, and (ii) the 25 external emetic ST 26 isolates (downloaded from RefSeq). ANOSIM tests were conducted using pairwise core SNP differences and 5 independent runs of 10,000 permutations each. For both the ANOVA-like permutation tests and the ANOSIM tests, Bonferroni corrections were used to correct for multiple comparisons at the α = 0.05 level.

### 2.14 Statistical comparison of phylogenetic trees

The Kendall-Colijn (Kendall and Colijn, 2015) test described by Katz, et al. (Katz et al., 2017) was used to compare the topologies of trees, using the treespace (Jombart et al., 2017), ips (Heibl, 2008), phangorn (Schliep, 2011), docopt (de Jonge, 2016), and stringr (Wickham, 2018) packages in R version 3.4.3. The phylogenies that underwent pairwise testing were constructed using core SNPs identified in (i) 30 emetic clade III genomes via all six SNP calling pipelines, and (ii) 55 emetic ST 26 genomes (25 publicly-available genomes and the 30 emetic isolates sequenced here) using the kSNP3 and Parsnp pipelines. For all pairwise tree comparisons, a lambda value of 0 was used along with 100,000 random trees as a background distribution, and a Bonferroni correction was used to correct for multiple comparisons. Two trees were considered to be more topologically similar than would be expected by chance if a significant *P* value (*P* < 0.05) resulted after correcting for multiple testing (Katz et al., 2017).

### 2.15 Calculation of average nucleotide identity values

FastANI version 1.0 (Jain, 2017) was used to calculate average nucleotide identity (ANI) values between assembled genomes of isolates sequenced in this study and selected reference genomes (Table 2), as well as the genomes of 18 currently-recognized *B. cereus* group species (Table 3).

**Table 3.** List of outbreak isolates and corresponding metadata, single- and multi-locus sequence types, and species.

| Isolate Name | Source (General) | Source (Specific) | Collection Date | Isolation Date | Production Date/Batch<sup>a</sup> | *panC* Clade<sup>b</sup> | MLST ST<sup>c</sup> | *rpoB* AT<sup>d</sup> | Closest Type Strain (ANI)<sup>e</sup> |
| --- | --- | --- | --- | --- | --- | --- | --- | --- | --- |
| FOOD_10_18_16_LFTOV_NA_R9-6400 | Food | Leftovers | 9-Oct-16 | 18-Oct-16 | Unknown | III | 26 | 125 | *B. paranthracis* MN5 (97.5) |
| FOOD_10_18_16_LFTOV_NA_R9-6401 | Food | Leftovers | 9-Oct-16 | 18-Oct-16 | Unknown | III | 26 | 125 | *B. paranthracis* MN5 (97.5) |
| FOOD_10_18_16_LFTOV_NA_R9-6402 | Food | Leftovers | 9-Oct-16 | 18-Oct-16 | Unknown | III | 26 | 125 | *B. paranthracis* MN5 (97.5) |
| FOOD_10_19_16_RSNT1_1B_R9-6388 | Food | Restaurant 1 | 6-Oct-16 | 19-Oct-16 | 1/B | III | 26 | 125 | *B. paranthracis* MN5 (97.5) |
| FOOD_10_19_16_RSNT1_1B_R9-6389 | Food | Restaurant 1 | 6-Oct-16 | 19-Oct-16 | 1/B | III | 26 | 125 | *B. paranthracis* MN5 (97.5) |
| FOOD_10_19_16_RSNT1_1B_R9-6390 | Food | Restaurant 1 | 6-Oct-16 | 19-Oct-16 | 1/B | III | 26 | 125 | *B. paranthracis* MN5 (97.5) |
| FOOD_10_19_16_RSNT1_1B_R9-6391 | Food | Restaurant 1 | 6-Oct-16 | 19-Oct-16 | 1/B | III | 26 | 125 | *B. paranthracis* MN5 (97.5) |
| FOOD_10_19_16_RSNT1_2A_R9-6386 | Food | Restaurant 1 | 6-Oct-16 | 19-Oct-16 | 2/A | III | 26 | 125 | *B. paranthracis* MN5 (97.5) |
| FOOD_10_19_16_RSNT1_2A_R9-6387 | Food | Restaurant 1 | 6-Oct-16 | 19-Oct-16 | 2/A | III | 26 | 125 | *B. paranthracis* MN5 (97.5) |
| FOOD_10_19_16_RSNT1_2H_R9-6392 | Food | Restaurant 1 | 6-Oct-16 | 19-Oct-16 | 2/H | III | 26 | 125 | *B. paranthracis* MN5 (97.5) |
| FOOD_10_19_16_RSNT1_2H_R9-6393 | Food | Restaurant 1 | 6-Oct-16 | 19-Oct-16 | 2/H | III | 26 | 125 | *B. paranthracis* MN5 (97.5) |
| FOOD_10_19_16_RSNT1_2H_R9-6394 | Food | Restaurant 1 | 6-Oct-16 | 19-Oct-16 | 2/H | III | 26 | 125 | *B. paranthracis* MN5 (97.5) |
| FOOD_10_19_16_RSNT1_2H_R9-6395 | Food | Restaurant 1 | 6-Oct-16 | 19-Oct-16 | 2/H | III | 26 | 125 | *B. paranthracis* MN5 (97.5) |
| FOOD_10_19_16_RSNT1_2H_R9-6396 | Food | Restaurant 1 | 6-Oct-16 | 19-Oct-16 | 2/H | III | 26 | 125 | *B. paranthracis* MN5 (97.5) |
| FOOD_10_19_16_RSNT2_2A_R9-6397 | Food | Restaurant 2 | 6-Oct-16 | 19-Oct-16 | 2/A | III | 26 | 125 | *B. paranthracis* MN5 (97.5) |
| FOOD_10_19_16_RSNT2_2A_R9-6398 | Food | Restaurant 2 | 6-Oct-16 | 19-Oct-16 | 2/A | III | 26 | 125 | *B. paranthracis* MN5 (97.5) |
| FOOD_10_19_16_RSNT2_2A_R9-6399 | Food | Restaurant 2 | 6-Oct-16 | 19-Oct-16 | 2/A | III | 26 | 125 | *B. paranthracis* MN5 (97.6) |
| FOOD_10_19_16_RSNT3_1E_R9-6407 | Food | Restaurant 3 | 6-Oct-16 | 19-Oct-16 | 1/E | III | 26 | 125 | *B. paranthracis* MN5 (97.5) |
| FOOD_10_19_16_RSNT3_2A_R9-6403 | Food | Restaurant 3 | 6-Oct-16 | 19-Oct-16 | 2/A | III | 26 | 125 | *B. paranthracis* MN5 (97.5) |
| FOOD_10_19_16_RSNT3_2A_R9-6404 | Food | Restaurant 3 | 6-Oct-16 | 19-Oct-16 | 2/A | III | 26 | 125 | *B. paranthracis* MN5 (97.5) |
| FOOD_10_19_16_RSNT3_2A_R9-6405 | Food | Restaurant 3 | 6-Oct-16 | 19-Oct-16 | 2/A | III | 26 | 125 | *B. paranthracis* MN5 (97.5) |
| FOOD_10_19_16_RSNT4_2B_R9-6408 | Food | Restaurant 4 | 6-Oct-16 | 19-Oct-16 | 2/B | III | 26 | 125 | *B. paranthracis* MN5 (97.5) |
| FOOD_10_19_16_RSNT4_2B_R9-6409 | Food | Restaurant 4 | 6-Oct-16 | 19-Oct-16 | 2/B | III | 26 | 125 | *B. paranthracis* MN5 (97.5) |
| FOOD_10_19_16_RSNT5_1C_R9-6411 | Food | Restaurant 5 | 6-Oct-16 | 19-Oct-16 | 1/C | III | 26 | 125 | *B. paranthracis* MN5 (97.5) |
| HUMN_10_18_16_FECAL_NA_R9-6384 | Human | Feces | 7-Oct-16 | 18-Oct-16 | NA | III | 26 | 125 | *B. paranthracis* MN5 (97.6) |
| HUMN_10_18_16_FECAL_NA_R9-6385 | Human | Feces | 8-Oct-16 | 18-Oct-16 | NA | III | 26 | 125 | *B. paranthracis* MN5 (97.5) |
| HUMN_10_18_16_FECAL_NA_R9-6412 | Human | Feces | 8-Oct-16 | 18-Oct-16 | NA | III | 26 | 125 | *B. paranthracis* MN5 (97.5) |
| HUMN_10_19_16_FECAL_NA_R9-6381 | Human | Feces | 7-Oct-16 | 19-Oct-16 | NA | III | 26 | 125 | *B. paranthracis* MN5 (97.5) |
| HUMN_10_19_16_FECAL_NA_R9-6382 | Human | Feces | 7-Oct-16 | 19-Oct-16 | NA | III | 26 | 125 | *B. paranthracis* MN5 (97.5) |
| HUMN_10_19_16_FECAL_NA_R9-6383 | Human | Feces | 7-Oct-16 | 19-Oct-16 | NA | III | 26 | 125 | *B. paranthracis* MN5 (97.5) |
| FOOD_10_19_16_RSNT3_1E_R9-6406 | Food | Restaurant 3 | 6-Oct-16 | 19-Oct-16 | 1/E | IV | 24 | 92 | *B. cereus* ATCC 14579 (98.9) |
| FOOD_10_19_16_RSNT5_1C_R9-6410 | Food | Restaurant 5 | 6-Oct-16 | 19-Oct-16 | 1/C | IV | 24 | 92 | *B. cereus* ATCC 14579 (98.9) |
| HUMN_10_26_16_FECAL_NA_R9-6413 | Human | Feces | 8-Oct-16 | 26-Oct-16 | NA | IV | 142 | 92 | *B. cereus* ATCC 14579 (98.8) |

<sup>a</sup>Production date is designated by either 1 or 2; batch is one of A through H
<sup>b</sup>*panC* clade assigned *in silico* using BTypser 2.2.0
<sup>c</sup>Multi-locus sequence typing (MLST) sequence type (ST) assigned *in silico* using BTypser 2.2.0
<sup>d</sup>*rpoB* allelic type (AT) determined using Sanger sequencing and verified *in silico* using BTypser 2.2.0
<sup>e</sup>ANI, average nucleotide identity calculated using FastANI

### 2.16 Availability of Data

Trimmed Illumina reads for all 33 isolates sequenced in this study have been made publicly available (NCBI BioProject Accession PRJNA437714), with NCBI Biosample accession numbers for all isolates listed in Supplementary Table S1. All figures have been deposited in FigShare (DOI https://doi.org/10.6084/m9.figshare.7001525.v1), and records of all isolates are available in Food Microbe Tracker (Vangay et al., 2013).

## 3 Results

### 3.1 Both emetic and diarrheal symptoms were reported among cases associated with the *B. cereus* foodborne outbreak

Between September 30^th^ and October 6^th^, 2016, local health departments in upstate New York’s Niagara and Erie counties reported a total of 179 estimated foodborne illness cases among customers of a Mexican fast-food restaurant chain in eight towns/cities. Among these cases, laboratory results were available for ten cases. For seven of these cases, *B. cereus* group species were isolated from patient stool samples. While no deaths, hospitalizations, or emergency room visits were reported from 169 cases from which information was obtained, 4 resulted in a visit to a health care provider (not including emergency room visits). More than 2/3 of 179 cases were female (69%), and 61% of cases fell within the 20-74 age group. In 156 of 179 total cases (87%), refried beans had been consumed.

Of 169 cases from which information was obtained, 88% reported vomiting, and more than half reported nausea and abdominal cramps (95 and 65%, respectively). However, in addition to vomiting, 38% of cases reported also diarrhea. Additional symptoms reported included (i) weakness (43%), (ii) chills (40%), (iii) dehydration (35%), (iv) headache (28%), (v) myalgia (muscle ache/pain; 16%), (vi) fever (16%), (vii) sweating (16%), and (viii) sore throat (3%). The incubation period observed for all cases ranged from 0.25-24 h, with a median of 2 h. The duration of illness ranged from 0.25 to 144 h, with a median estimate of 6 h.

A traceback was conducted, with the source of the outbreak determined to be a processing plant in Pennsylvania. The distributor in Pennsylvania packaged the refried beans specifically for the chain establishment where the outbreak occurred. The establishments where the outbreak occurred received 5 lb trays of pre-cooked, sealed, and frozen refried beans from the production/packaging facility. The refried beans would undergo cooking and a hot hold prior to consumption at the establishments where the outbreak occurred. It was determined that the refried beans were contaminated prior to preparation at the chain establishment.

Stool samples from suspect cases were cultured on MYP agar and *B. cereus*-like colonies were isolated from seven stool samples. Additionally, *B. cereus-*like colonies were isolated from nine food samples that were collected from five restaurants. In total, seven isolates from stool samples and 26 isolates from foods were confirmed to belong to the *B. cereus* group using standard microbiological methods. Isolates that were large Gram-positive rods, beta-hemolytic, and motile were presumptively identified as *B. cereus*-like. Additionally, spore staining was performed to differentiate between *B. cereus* and *B. thuringiensis.* All isolates were negative for the presence of parasporal crystals, therefore the isolates were classified as *B. cereus.* All 33 *B. cereus* group isolates underwent preliminary molecular characterization by Sanger sequencing of *rpoB*, which revealed two distinct allelic types belonging to phylogenetic clades III (*rpoB* allelic type 125; AT 125) and IV (AT 92).

### 3.2 WGS confirms presence of multiple *B. cereus* group species represented among strains sequenced in association with the outbreak

*rpoB* allelic types (ATs) assigned *in silico* were identical to those obtained using Sanger sequencing for all 33 isolates (Table 3). *panC* clade assignment confirmed the presence of *B. cereus* from multiple clades (Table 3), with clade III (n = 30) and clade IV (n = 3) represented among the 33 isolates. *In silico* MLST further resolved the clade IV isolates into two sequence types (STs): the two strains isolated from refried beans served at two different restaurants had identical STs, while the single human isolate belonging to clade IV had a unique ST (Table 3). All 30 *panC* clade III isolates belonged to ST 26, including the remaining six human clinical isolates (Table 3).

The presence of isolates from multiple *B. cereus* group clades, as suggested by the *rpoB, panC,* and MLST loci among isolates sequenced in conjunction with this outbreak was confirmed using core SNPs detected in all outbreak isolates, as well as the genomes of 18 currently-recognized *B. cereus* group species (Figure 1). The three isolates assigned to *panC* clade IV using a 7-clade scheme (Guinebretiere et al., 2008) were most closely related to the *B. cereus s.s.* type strain (Figure 1). All three clade IV *B. cereus* isolates possessed diarrheal toxin genes *hblABCD* and *cytK2* at high identity and coverage (Figure 1), which code for enterotoxins hemolysin BL (Hbl) and cytotoxin K (CytK), respectively. The 30 isolates assigned to *panC* clade III, however, were most closely related to the type strain of *B. paranthracis* (Figure 1). Unlike *B. paranthracis,* all of the clade III isolates investigated here possessed the *cesABCD* operon (Figure 1), which codes for emetic toxin-producing cereulide synthetase (in the case of isolate HUMN_10_18_16_FECAL_NA_R9-6384, *cesD* was split onto two contigs), and were motile.

**Figure 1.**
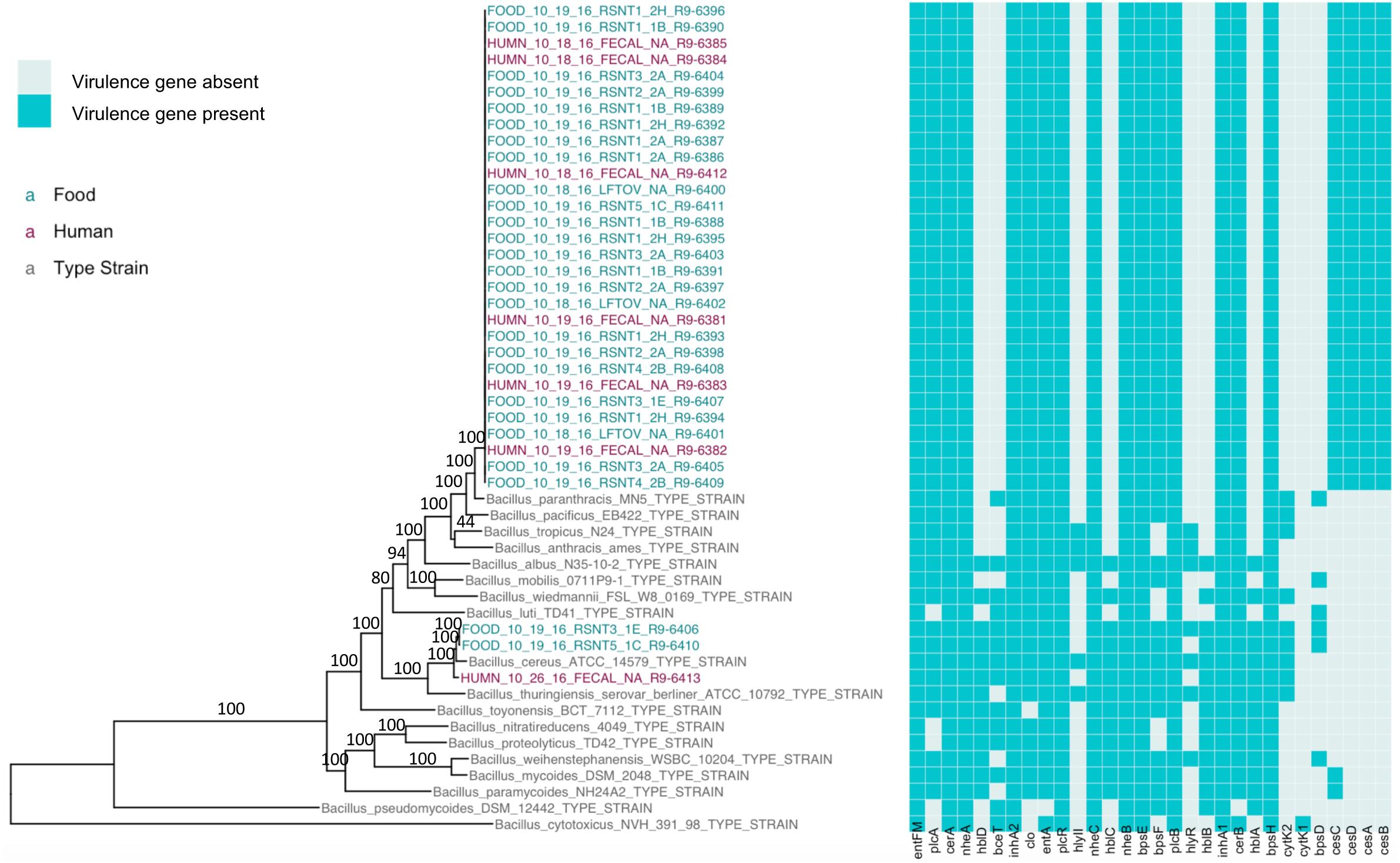
Maximum likelihood phylogeny of core SNPs identified in 33 isolates sequenced in conjunction with a *B. cereus* outbreak, as well as genomes of the 18 currently recognized *B. cereus* group species (shown in gray). Core SNPs were identified in all genomes using kSNP3. Heatmap corresponds to presence/absence of *B. cereus* group virulence genes detected in each sequence using BTyper. Tip labels in maroon and teal correspond to the 7 human clinical isolates and 26 isolates from food sequenced in conjunction with this outbreak, respectively. Phylogeny is rooted at the midpoint, and branch labels correspond to bootstrap support percentages out of 500 replicates.

Based on average nucleotide identity (ANI) values, the three diarrheal clade IV isolates were classified as *B. cereus s.s.* (ANI > 95; Table 3). The 30 emetic clade III isolates, however, did not meet the minimum ANI cutoff of 95 used for assigning bacterial species relative to the *B. cereus s.s.* type strain. Of the 18 *B. cereus* group species as they are currently defined (Liu et al., 2017), the *B. paranthracis* type strain was closest to the 30 emetic clade III isolates from this outbreak (ANI > 95; Table 3), indicating that the emetic clade III and diarrheal clade IV isolates from this outbreak are different *B. cereus* group species.

### 3.3 Emetic and diarrheal *B. cereus* isolates associated with the foodborne outbreak do not differ in cytotoxicity

All three diarrheal strains isolated in conjunction with the outbreak (FSL R9-6406, FSL R9-6410, and FSL R9-6413) were found to produce Hbl, as well as non-hemolytic enterotoxin (Nhe). Characterization of six representatives of the emetic isolates tested (i.e., FSL R9-6381, FSL R9-6382, FSL R9-6384, FSL R9-6389, FSL R9-6395, and FSL R9-6399) revealed that they produced Nhe, but not Hbl. Supernatants of diarrheal *B. cereus s.s.* ATCC 14579 showed stronger inhibitory effect on the viability of HeLa cells compared to supernatants of the 33 outbreak-associated isolates (*P* < 0.05; Figure 2). Furthermore, the viability of HeLa cells treated with 0.05% Triton X-100, the positive control, was significantly lower compared to viability of HeLa cells treated with bacterial supernatants (Games-Howell *P* < 0.05; Figure 2). Among all pairs of emetic isolates, only the viabilities of HeLa cells exposed to the supernatants of isolates FSL R9-6409 and FSL R9-6387 were found to differ (*P* < 0.05; Figure 2). The differences in HeLa cell viability after treatment with supernatants of these two emetic outbreak-associated strains are likely due to biological variability among replicates, as outbreak-associated emetic isolates were shown to be clonal (Figure 1). Taken together, the emetic group (represented by 30 emetic outbreak-associated isolates) had a mean cell viability of 97.5 ± 5.1%, while the diarrheal group (represented by 3 diarrheal outbreak-associated isolates) gave a mean cell viability of 101.4 ± 7.9%.

**Figure 2.**
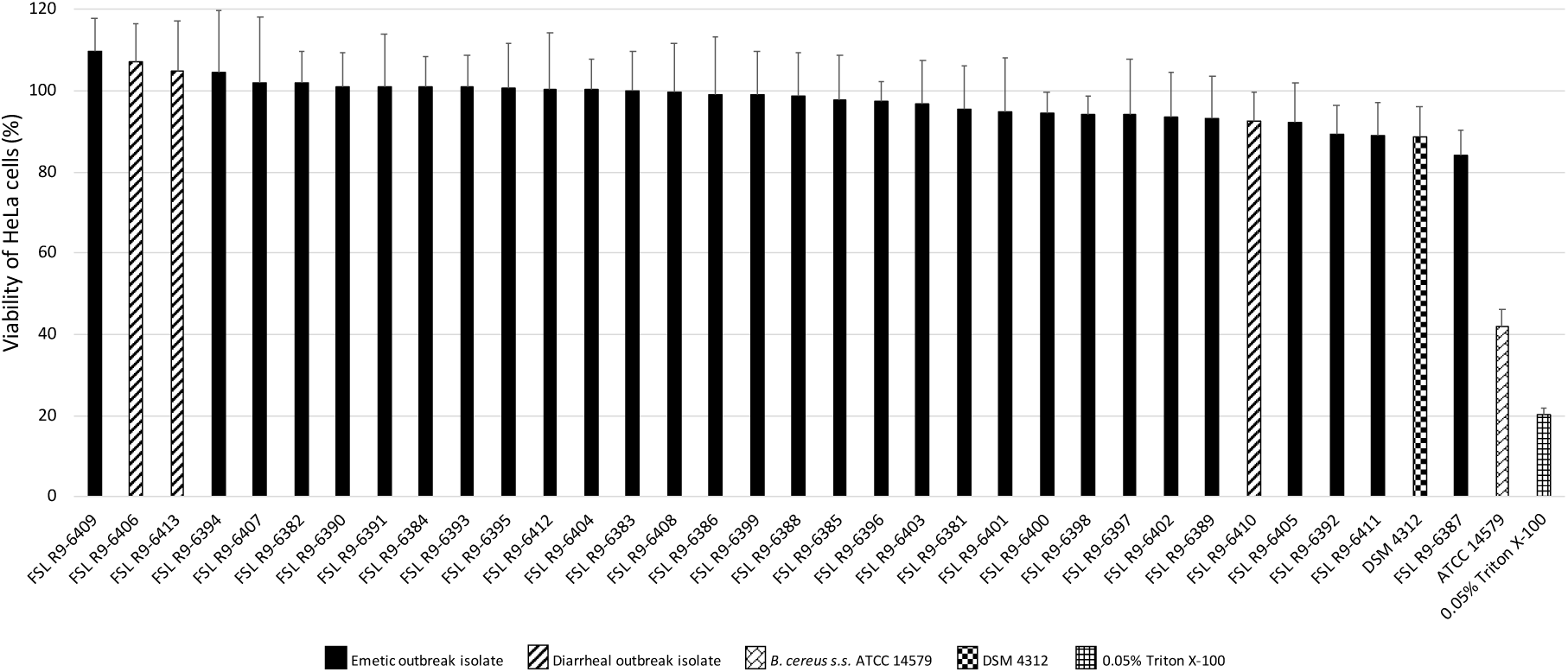
Percentage viability of HeLa cells when treated with supernatants of each isolate as determined by the WST-1 assay. Viability was calculated as ratio of corrected absorbance of solution when HeLa cells were treated with supernatants to the ratio of corrected absorbance of solution when HeLa cells were treated with BHI (i.e., negative control), converted to percentages. The columns represent the mean viabilities, while the error bars represent standard deviations for 12 technical replicates.

### 3.4 Core SNPs identified among *B. cereus* group outbreak isolates from two clades are dependent on variant calling pipeline and reference genome selection

To simulate a scenario in which genomes from a *B. cereus* outbreak spanning multiple clades were analyzed in aggregate, core SNPs were identified in all 33 outbreak isolates from clades III and IV (n = 30 and 3 isolates, respectively) using (i) combinations of five reference-based variant calling pipelines (Table 1) and three different reference genomes (Table 2) and (ii) a reference-free SNP calling method (Table 1). When genomes from all 33 isolates were analyzed together, the numbers of SNPs identified by each pipeline and reference combination varied by up to several orders of magnitude (Figure 3A), often with little agreement between pipelines in terms of the SNPs they reported (Figure 4). Independent of reference genome, the CFSAN pipeline was the most conservative, consistently identifying the fewest number of core SNPs when all 33 isolates were queried in aggregate (50, 27, and 0 core SNPs using reference genomes from clade III, IV, and VII, respectively) (Figure 3A). This can be contrasted with the Samtools, Freebayes, and Parsnp pipelines, which produced upwards of 100,000 core SNPs when the selected reference genome was a member of one of the clades being queried in the outbreak isolate set (clade III and IV; Figure 3A). In cases where a distant genome was used as the reference (clade VII’s *B. cytotoxicus* type strain chromosome), all reference-based pipelines reported fewer core SNPs than kSNP3’s reference-free *k-*mer based SNP calling approach (Figure 3A).

**Figure 3.**
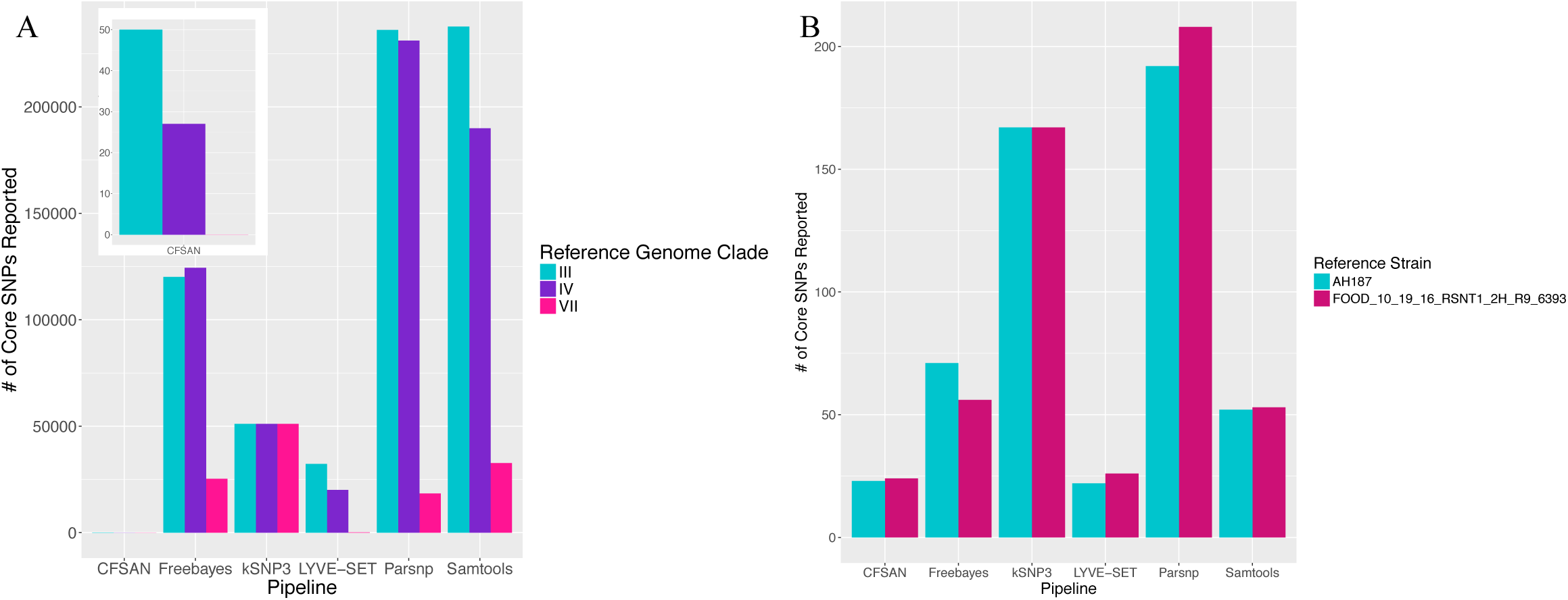
Number of core SNPs identified in (A) 33 *B. cereus* group isolates from two clades (30 and 3 isolates from clades III and IV, respectively) and (B) 30 emetic *B. cereus* group isolates from clade III, sequenced in conjunction with a foodborne outbreak. Combinations of five reference-based variant calling pipelines and (A) three and (B) two reference genomes, as well as one reference-free SNP calling method (kSNP3), were tested.

**Figure 4.**
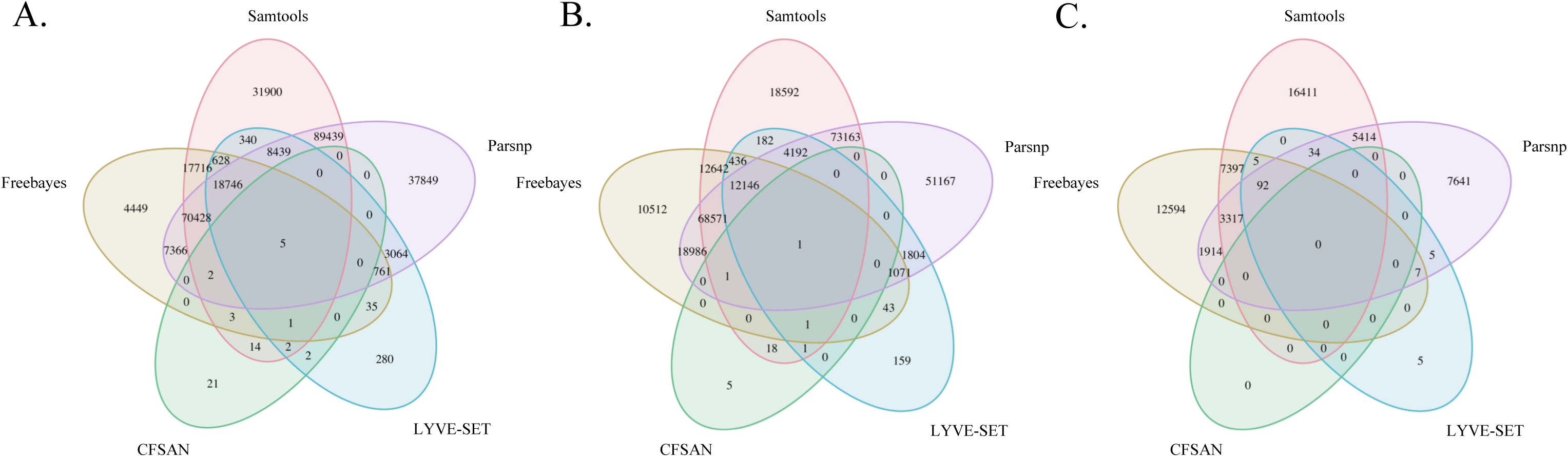
Comparison of SNP positions reported by five variant-calling pipelines for 33 *B. cereus* group strains isolated in association with a foodborne outbreak, with the chromosomes of (A) *B. cereus* str. AH187 (Clade III), (B) *B. cereus s.s.* str. ATCC 14579 (Clade IV), and (C) *B. cytotoxicus* str. NVH 391-98 (Clade VII) used as reference genomes. Ellipses represent each pipeline.

### 3.5 Choice of variant calling pipeline has greater influence on core SNP identification than choice of closely-related closed or draft reference genome for emetic clade III *B. cereus* group isolates

The 30 emetic clade III isolates were queried in the absence of their clade IV counterparts using combinations of five reference-based variant calling pipelines (Table 1) and two reference genomes (the closed chromosome of *B. cereus* str. AH187 and contigs of one of the isolates identified in this outbreak; Table 2) and one reference-free SNP calling method (Table 1). In this scenario, the choice of variant calling pipeline had a greater effect on the number of core SNPs obtained than the choice of reference genome, as both reference genomes possessed the same virulence gene profile (virulotype), *rpoB* AT, *panC* clade, MLST sequence type, and were of the same species (*B. paranthrasis* ANI > 95) as the 30 emetic isolates (Figure 3B). Congruent with this, the number of pairwise core SNP differences between emetic isolates sequenced in this outbreak varied more with the selection of variant calling pipeline than with reference genome (Figure 5). When the closed chromosome of *B. cereus* str. AH187 was used as a reference, pairwise core SNP differences among emetic isolates from this outbreak ranged from 0 to 8 (mean of 2.9; CFSAN), 7 to 29 (mean of 16.1; Freebayes), 0 to 8 (mean of 2.8; LYVE-SET), 0 to 64 (mean of 23.6; Parsnp), and 1 to 16 SNPs (mean of 8.2; Samtools) (Figure 5). Using the reference-free kSNP3 pipeline, this range was 1 to 46 SNPs (mean of 16.7; Figure 5). The CFSAN and LYVE-SET pipelines produced nearly identical results in terms of the number and identity of the core SNPs called (23 and 22 SNPs, respectively; Figure 6), while the two methods that relied on assembled genomes rather than short reads for SNP calling (kSNP3 and Parsnp) produced the greatest numbers of core SNPs (Figure 3B). The topologies of phylogenies constructed using core SNPs identified by each of the six pipelines also reflected this, as the topologies of the CFSAN/LYVE-SET and kSNP3/Parsnp pipelines were more similar to each other than what would be expected by chance (Table 4 and Figure 7).

**Table 4.**
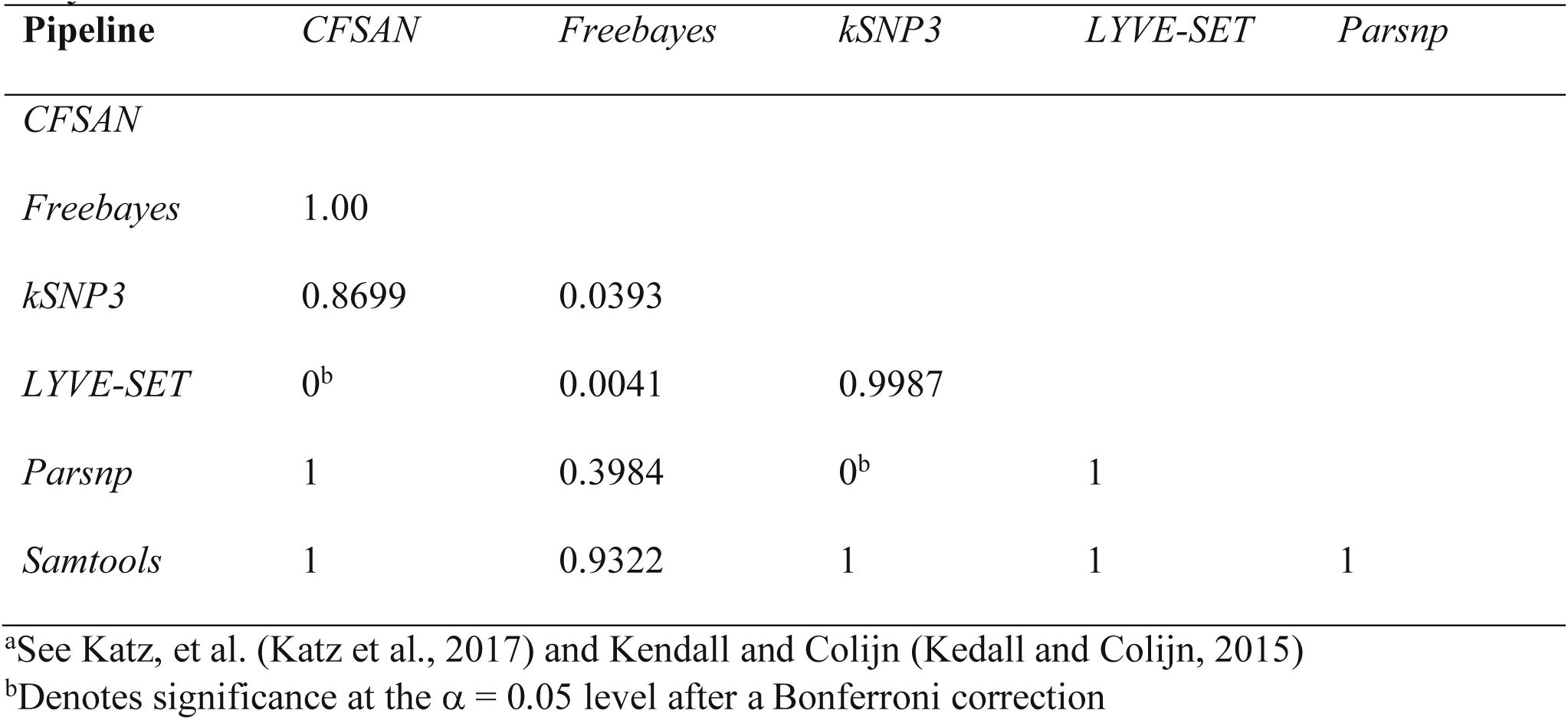
*P-*values obtained from pairwise tests of tree topologies using a *Z* test based on the Kendall-Colijn metric.^a^

| Pipeline | *CFSAN* | *Freebayes* | *kSNP3* | *LYVE-SET* | *Parsnp* |
| --- | --- | --- | --- | --- | --- |
| *CFSAN* |  |  |  |  |  |
| *Freebayes* | 1.00 |  |  |  |  |
| *kSNP3* | 0.8699 | 0.0393 |  |  |  |
| *LYVE-SET* | 0<sup>b</sup> | 0.0041 | 0.9987 |  |  |
| *Parsnp* | 1 | 0.3984 | 0<sup>b</sup> | 1 |  |
| *Samtools* | 1 | 0.9322 | 1 | 1 | 1 |

<sup>a</sup>See Katz, et al. (Katz et al., 2017) and Kendall and Colijn (Kedall and Colijn, 2015)
<sup>b</sup>Denotes significance at the $\alpha = 0.05$ level after a Bonferroni correction

**Figure 5.**
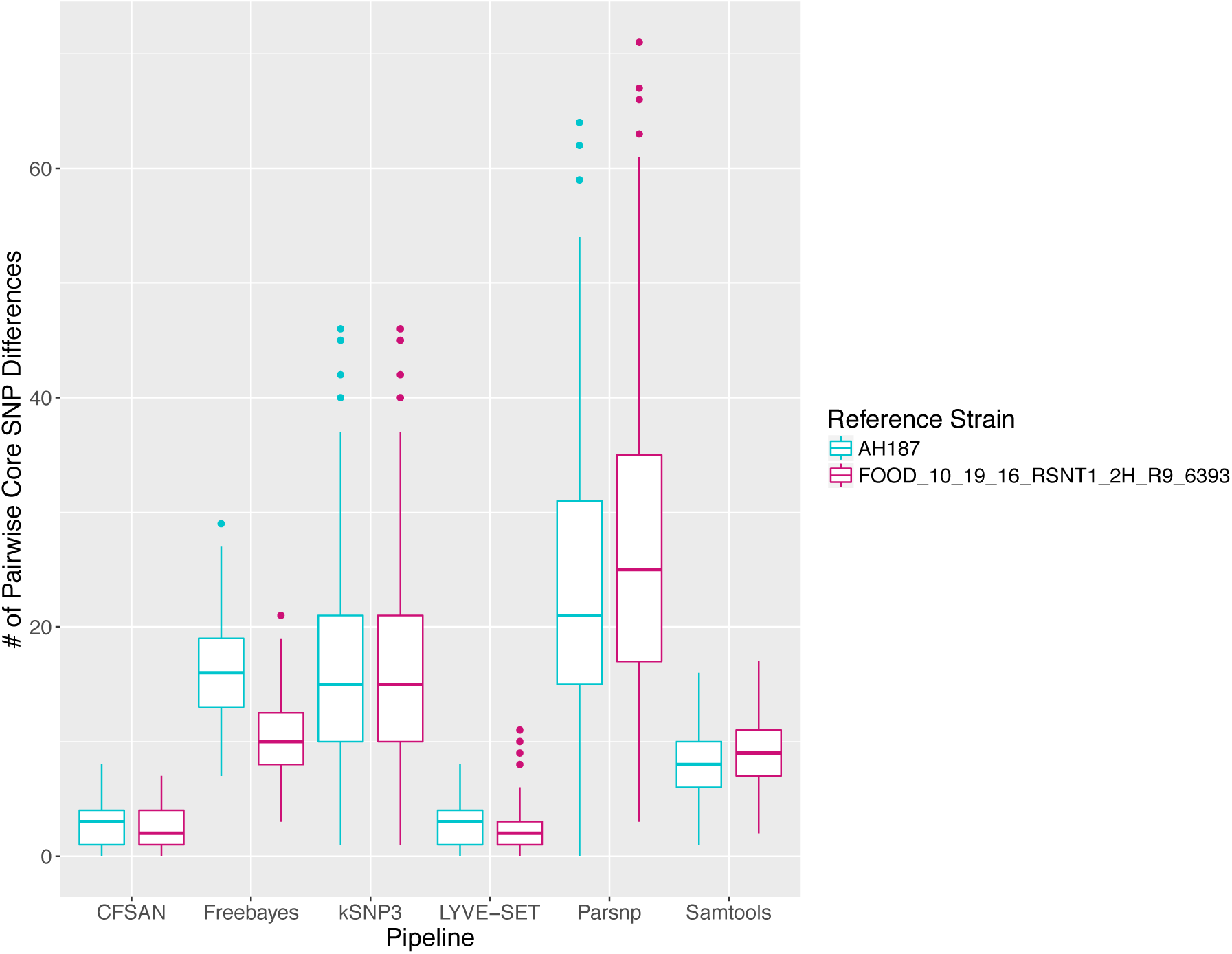
Ranges of pairwise core SNP differences between 30 emetic clade III *B. cereus* group strains isolated in conjunction with a foodborne outbreak. Combinations of five reference-based variant calling pipelines and two reference genomes, as well as one reference-free SNP calling method (kSNP3) were tested. Lower and upper box hinges correspond to the first and third quartiles, respectively. Lower and upper whiskers extend from the hinge to the smallest and largest values no more distant than 1.5 times the interquartile range from the hinge, respectively. Points represent pairwise distances that fall beyond the ends of the whiskers.

**Figure 6.**
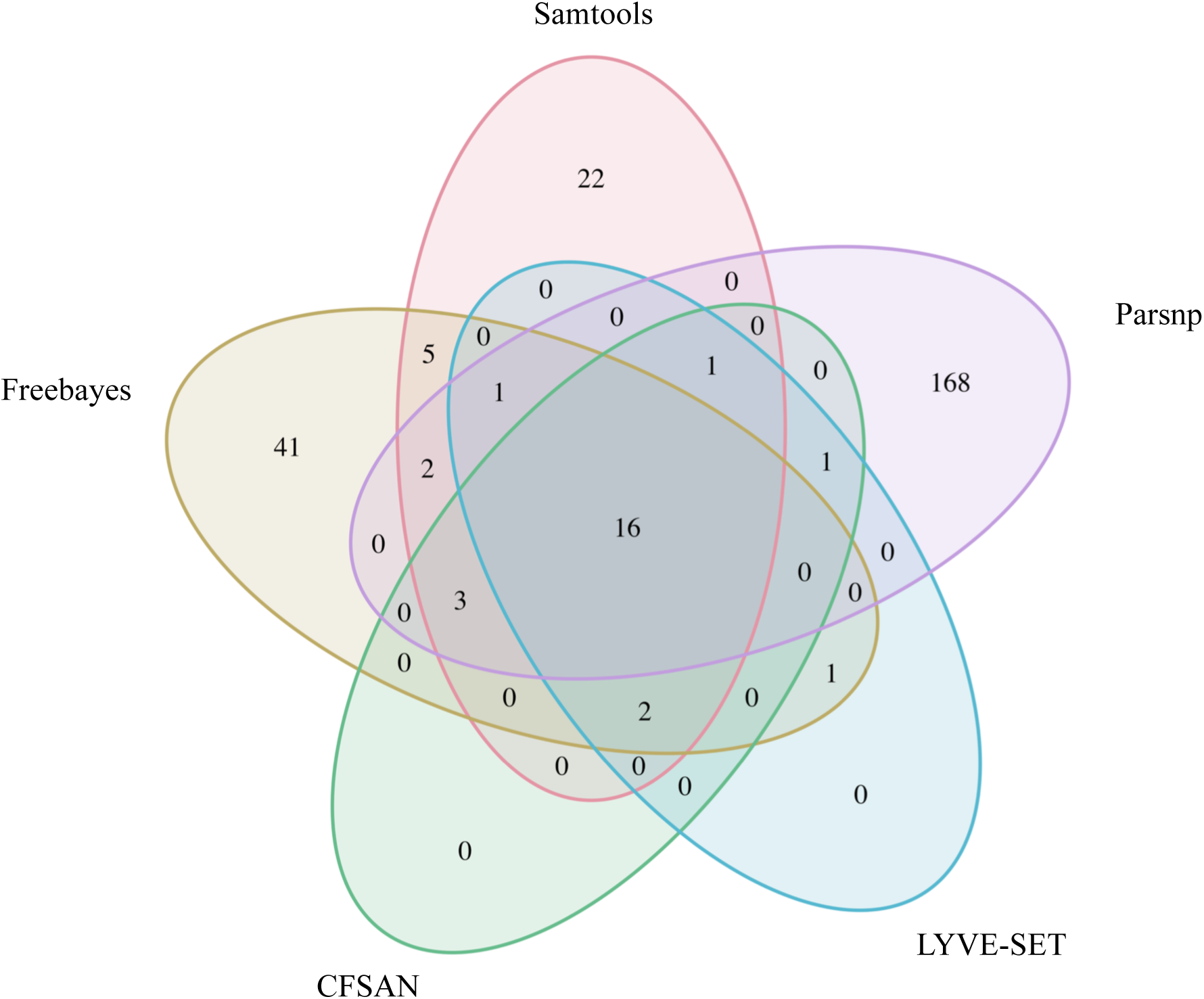
Comparison of SNP positions reported by five variant-calling pipelines for 30 emetic clade III *B. cereus* group outbreak isolates. Ellipses represent each pipeline, all of which used the chromosome of emetic clade III *B. cereus* strain AH187 as a reference for variant calling.

**Figure 7.**
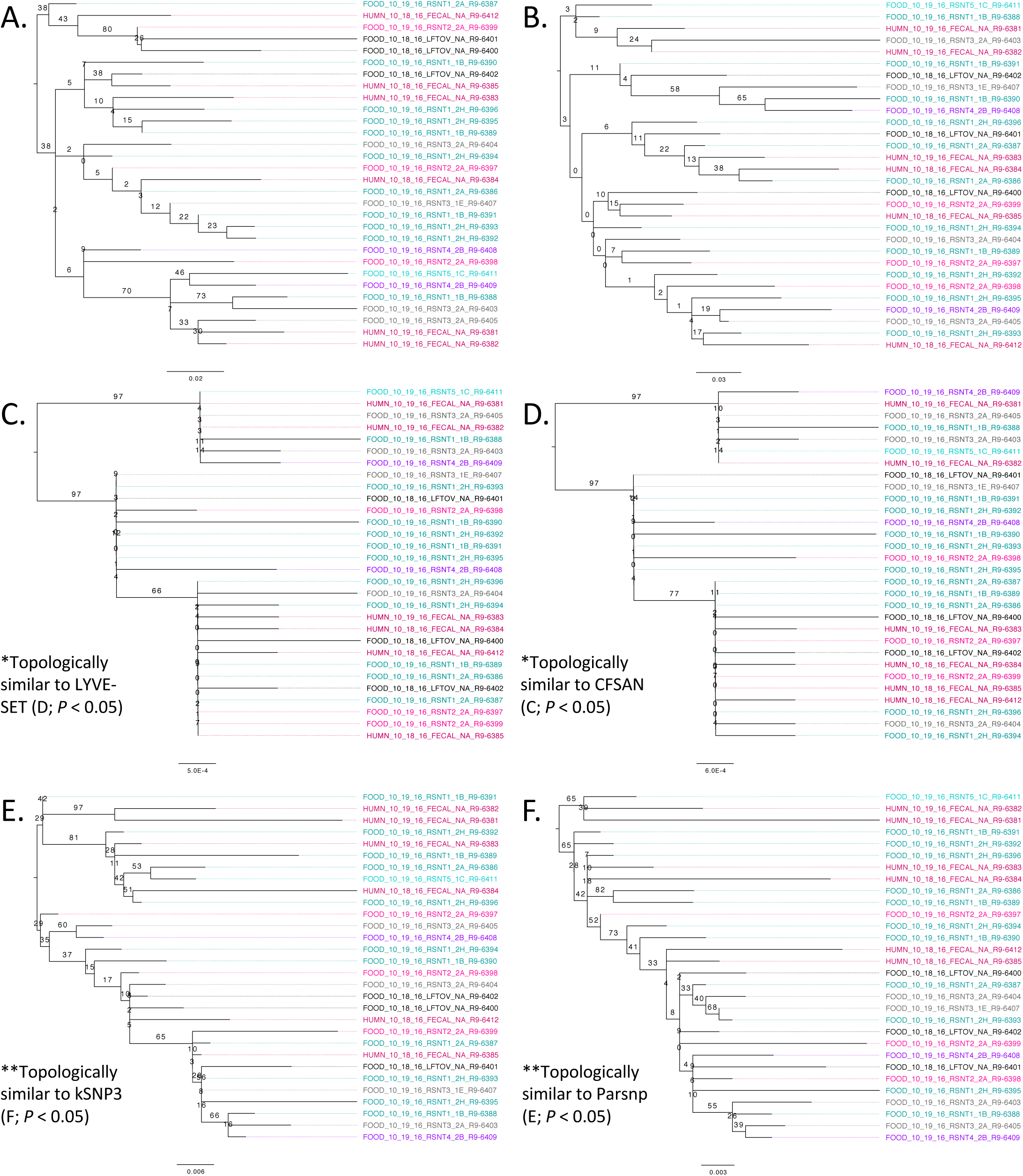
Maximum likelihood phylogenies constructed using core SNPs detected in 30 emetic clade III outbreak isolates using the (A) Samtools, (B) Freebayes, (C) CFSAN, (D) LYVE-SET, (E) Parsnp, and (F) kSNP3 variant calling pipelines using *B. cereus* str. AH187 as reference. Branch labels correspond to bootstrap support percentages out of 1,000 replicates, while like-colored tip labels correspond to isolates from the same source (human clinical fecal sample, leftovers, or restaurant 1, 2, 3, 4, or 5).

Within the emetic clade III isolates associated with this outbreak, a total of 32 core SNPs were identified by two or more of the reference-based variant calling pipelines when *B. cereus* str. AH187 was used as a reference, half of which were identified by all 5 pipelines (Figure 6). Out of these 32 SNPs, 23 were identified in protein coding genes, 14 of which produced non-synonymous amino acid changes (Supplementary Table S3). Genes with non-synonymous changes were involved in molybdopterin biosynthesis (WP_000544623.1), proteolysis (WP_000215096.1 and WP_000857793.1), chitin binding (WP_000795732.1), iron-hydroxamate transport (WP_000728195.1), DNA repair (WP_000947749.1 and WP_000867556.1), DNA replication (WP_000867556.1 and WP_000435993.1), protein transport and insertion into the membrane (WP_000727745.1), and glyoxylase/bleomycin resistance (WP_000800664.1).

### 3.6 Phylogenies constructed using core SNPs identified in 55 emetic ST 26 *B. cereus* isolates by kSNP3 and Parsnp yield similar topologies

To compare the 30 emetic strains from this outbreak to other emetic clade III isolates, all emetic clade III genomes with ST 26 were downloaded from NCBI. This produced a total of 55 emetic clade III isolates with ST 26 (30 isolates from this outbreak plus 25 from NCBI RefSeq). Among the 55 emetic ST 26 genomes, Parsnp identified almost twice as many core SNPs as kSNP3 (4,597 and 2,593 core SNPs, respectively). However, the topologies of phylogenies produced using the core SNPs identified by each pipeline were found to be more similar than would be expected by chance (Kendall-Colijn test *P <* 0.05; Figure 8).

**Figure 8.**
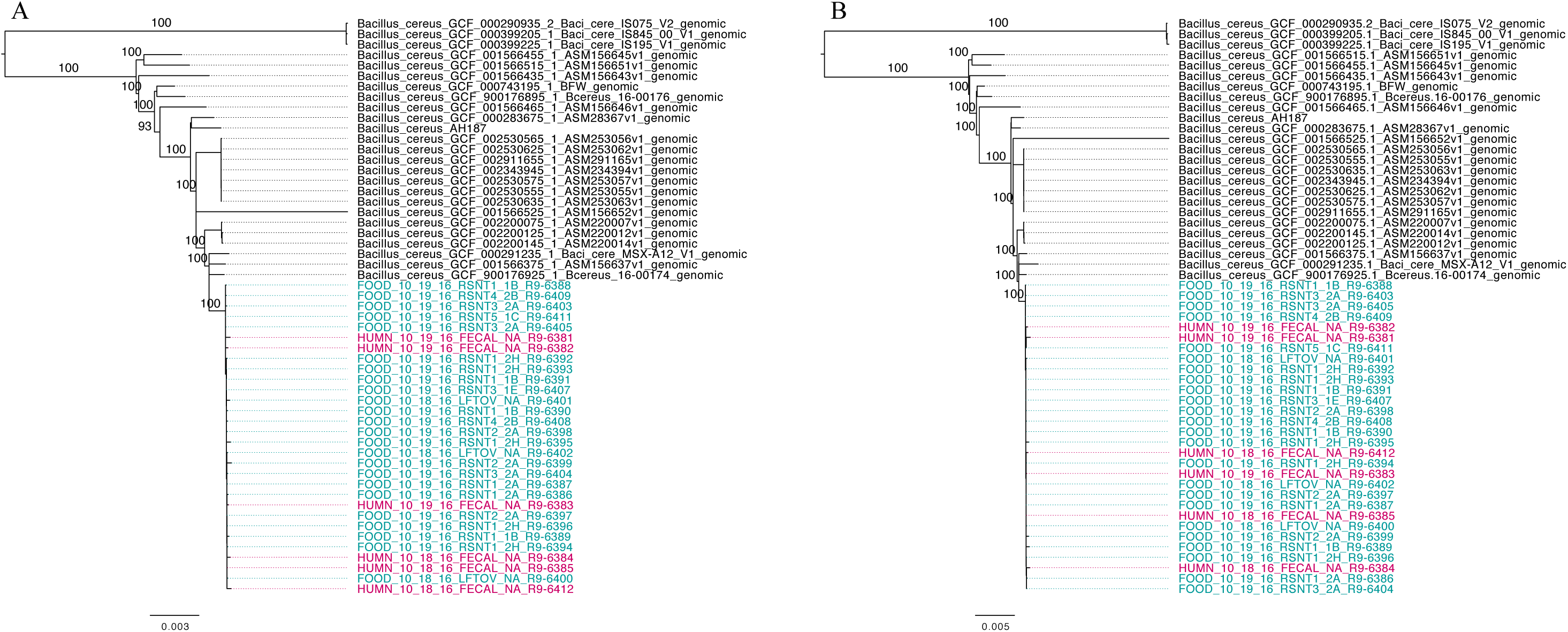
Maximum likelihood phylogenies of 30 emetic clade III isolates (ST 26) sequenced in conjunction with a *B. cereus* outbreak, as well as all other emetic clade III ST 26 genomes available in NCBI (n = 25; shown in black). Trees were constructed using core SNPs identified using (A) kSNP3 or (B) Parsnp. Tip labels in maroon and teal correspond to the 6 human clinical isolates and 24 isolates from food sequenced in conjunction with this outbreak, respectively. Branch labels correspond to bootstrap support percentages out of 1,000 replicates.

Based on pairwise core SNP differences, the publicly-available genomes showed greater variability than the outbreak isolates described here, regardless of whether kSNP3 or Parsnp was used for variant calling (ANOVA-like permutation test *P <* 0.05). Pairwise core SNP differences of the 30 emetic clade III isolates from this outbreak ranged from 0 to 25 SNPs (mean of 8.3) and 0 to 44 SNPs (mean of 11.9) when the kSNP3 and Parsnp pipelines were used, respectively. For external ST 26 isolates not associated with this outbreak, pairwise core SNP differences ranged from 0 to 1,474 SNPs (mean of 425.7) and 0 to 3,111 SNPs (mean of 828.3) when kSNP3 and Parsnp were used, respectively. Between these two groups (the 30 emetic isolates from this outbreak and the 25 external emetic ST 26 isolates), pairwise core SNP differences ranged from 73 to 1,258 SNPs (mean of 301.7; kSNP3) and 74 to 2,709 SNPs (mean of 528.0; Parsnp). Reflecting this, the average of the ranks of pairwise SNP distances within emetic isolates from this outbreak was less than the average of the ranks of pairwise SNP distances between the emetic isolates from this outbreak and the external ST 26 isolates (ANOSIM *P* < 0.05). This is likely a result of the differences in variance between the outbreak and external ST 26 isolates, as supported by the results of the ANOVA-like permutation test (Anderson and Walsh, 2013).

## 4. Discussion

While *B. cereus* causes a considerable number of foodborne illnesses cases annually, outbreaks are rarely investigated with the methodological vigor (e.g., use of WGS) that is increasingly used for surveillance and outbreak investigations targeting other foodborne pathogens. A specific challenge in the U.S. is that, unlike for some other diseases, disease cases caused by *B. cereus* are typically not reportable, even though foodborne illnesses, regardless of etiology, are reportable in some states, including NY. This, combined with the typically mild course of *B. cereus* infection, means that human *B. cereus* isolates are rarely available for WGS. Furthermore, even if clinical *B. cereus* group isolates are available, WGS may not be used for isolate characterization in cases where infections are mild. Due to the availability of *B. cereus* isolates for seven human cases, the outbreak reported here presented a unique opportunity to pilot the use of WGS for investigation of *B. cereus* outbreaks. The data and approaches presented here will not only facilitate future investigation of other *B. cereus* outbreaks, but will also help with application of WGS for investigation of other foodborne disease outbreaks where limited reference WGS data and information on genomic diversity are available.

### 4.1 Considerations for addressing the unique challenges associated with characterization of foodborne outbreaks linked to the *B. cereus* group

In *B. cereus* outbreaks, interpretation of WGS data can be challenging, especially in cases where strains of multiple closely related species or subtypes appear to be associated with an outbreak. *B. cereus* outbreaks—particularly emetic outbreaks caused by cereulide-producing *B. cereus* group isolates—are often associated with improper handling of food (e.g., temperature abuse) (Ehling-Schulz et al., 2004; Stenfors Arnesen et al., 2008). This, and their ubiquitous presence in the environment, make it important to consider the possibility of a multi-strain or multi-species outbreak in addition to a single-source outbreak caused by a single strain. In the outbreak characterized here, *B. cereus* group strains from two phylogenetic clades, III and IV, were isolated from both human clinical stool samples, as well as refried beans from food samples linked to the outbreak. The separation of outbreak-related isolates into three diarrheal clade IV isolates (representing two distinct STs) and 30 emetic isolates may be explained by one of the following scenarios: (i) the outbreak was caused by refried beans contaminated with multiple *B. cereus* group species (isolates from clades III and IV), both of which caused illness in humans, (ii) in addition to housing emetic outbreak strains that belonged to clade III, samples of refried beans and patient stool samples harbored clade IV *B. cereus* group isolates that were not part of the outbreak but were incidentally isolated from stool and food samples, or (iii) a subset of patient stool samples and food samples did not harbor *B. cereus* group clade III isolates belonging to the outbreak, but did harbor clade IV strains that were isolated and sequenced. In order to determine which of these scenarios explains the presence of multiple *B. cereus* species among isolates sequenced in conjunction with a foodborne outbreak, additional epidemiological and microbiological data are needed.

Valuable metrics for inclusion/exclusion of *B. cereus* group cases in a foodborne outbreak include patient exposure, patient symptoms (e.g., vomiting, diarrhea, onset and duration of illness), levels of *B. cereus* present in implicated food and patient samples (CFU/g or CFU/ml), cytotoxicity of isolates, and the approach used to select bacterial colonies to undergo WGS (e.g., Glasset et al., 2016 recommend collecting at least five colonies representing a range of morphologies from each potentially contaminated food sample). However, some of these data may be more valuable than others: in their characterization of 564 *B. cereus* group strains associated with 140 “strong-evidence” foodborne outbreaks in France between 2007 and 2014, Glasset, et al. (Glasset et al., 2016) found that patient symptoms could not be associated with the presence of emetic and diarrheal strains. More than half (57%) of the *B. cereus* outbreaks queried in their study included patients exhibiting both emetic and diarrheal symptoms. Similar results were observed here, as emetic and diarrheal symptoms were reported in 88 and 38% of cases, respectively, with both vomiting and diarrhea reported by multiple patients. While it has been proposed that this may be due to the fact that emetic clade III isolates have been shown to produce diarrheal enterotoxin Nhe at high levels (Glasset et al., 2016), incongruences between isolate virulotype and patient symptoms may still exist.

Another metric that can be used for determining whether *B. cereus* group isolates are part of an outbreak or not is the level of *B. cereus* present in the implicated food. Like patient symptoms, *B. cereus* counts from implicated foods may aid in an outbreak investigation, but likely cannot definitively prove whether an isolate is part of an outbreak or not. For example, outbreaks caused by implicated foods with *B. cereus* counts of < 10^3^ CFU/g and as low as 400 CFU/g for diarrheal and emetic diseases, respectively, have been described (Glasset et al., 2016), despite levels of at least 10^5^/g being often detected in implicated foods (Stenfors Arnesen et al., 2008). The levels of *B. cereus* present in refried beans in the outbreak described here were not determined. However, like patient symptoms, *B. cereus* count data may be a useful supplemental metric for characterizing outbreak isolates in the future.

Incubation period can also be used to determine whether an isolate is part of an outbreak or not, as it is significantly shorter for emetic strains than diarrheal strains (Ehling-Schulz et al., 2004; Stenfors Arnesen et al., 2008; Glasset et al., 2016). In the outbreak described here, the patient from which a non-emetic clade IV *B. cereus* group strain was isolated reported an incubation time of 1 h, the lowest incubation time of all seven confirmed human clinical cases. However, this is still within the observed range of incubation times for emetic *B. cereus* disease (0.5 – 6 h) (Stenfors Arnesen et al., 2008), making it possible that the patient could have been infected with either emetic *B. cereus* that was part of the outbreak but not isolated, or a pathogen which caused similar symptoms to foodborne illness caused by emetic *B. cereus*.

Cytotoxicity data may also be leveraged to include/exclude outbreak-associated *B. cereus* group isolates. In the outbreak described here, the patient from which a non-emetic clade IV *B. cereus* group strain was isolated reported vomiting and nausea and no diarrheal symptoms, despite the clinical isolate’s possession of multiple diarrheal toxin genes and no emetic toxin genes. This could suggest that the *B. cereus* group strain isolated from the patient was not responsible for the illness but may also indicate that our understanding of the specific virulence genes responsible for different *B. cereus-*associated disease symptoms is still incomplete. To further investigate this, we carried out immunoassay-based detection of Hbl and Nhe, as well as a WST-1 proliferation assay on HeLa cells exposed to bacterial supernatants presumably containing toxins. The results of Hbl and Nhe immunodetection and cytotoxicity revealed that diarrheal isolates only had mild detrimental effects on HeLa cell viability, despite the fact that they produced hemolysin BL and nonhemolytic enterotoxin. This can be contrasted with the *B. cereus s.s.* type strain, which substantially reduced the viability of the HeLa cells.

For the outbreak described here, results obtained using a combination of microbiological, epidemiological, and bioinformatic methods indicate that hypothesis (i), in which the diarrheal strains were part of a multi-species outbreak, can likely be excluded. Evidence supporting the conclusion that the human clinical diarrheal isolate was not part of the outbreak described here include: (i) the emetic symptoms reported by the patient were incongruent with the virulotype of the isolate, (ii) the isolate had a different ST compared to all other isolates sequenced in this outbreak, and (iii) the isolate did not exhibit substantial cytotoxicity against HeLa cells (Figure 2). This may be due to the fact that this case was not part of the outbreak and was due to an infection or intoxication caused by another pathogen that leads to disease symptoms similar to *B. cereus* (e.g., *Staphylococcus aureus*), or that this person was an asymptomatic carrier of clade IV *B. cereus* (Ghosh, 1978; Turnbull and Kramer, 1985) that was isolated and sequenced instead of the clade III emetic outbreak isolate.

While we have shown here that WGS data can be a valuable tool for characterizing *B. cereus* group isolates from a foodborne outbreak, our results also showcase the importance of supplementing WGS data with epidemiological metadata to draw meaningful conclusions from *B. cereus* group genomic data. Furthermore, the availability of WGS and cytotoxicity data from a larger set of *B. cereus* isolates from symptomatic patients may also provide an opportunity to use comparative genomics approaches to further explore virulence genes that are linked to different disease outcomes in the future.

### 4.2 Recommendations for analyzing Illumina WGS data from *B. cereus* group isolates potentially linked to a foodborne outbreak

WGS is being used increasingly to characterize isolates associated with foodborne disease cases and outbreaks, and rightfully so: it offers the ability to characterize foodborne pathogens at unprecedented resolution, and it has been able to improve outbreak and cluster detection for numerous foodborne pathogens (Allard et al., 2017; Kovac et al., 2017; Moran-Gilad, 2017; Taboada et al., 2017), including *Salmonella enterica* (Taylor et al., 2015; Hoffmann et al., 2016; Gymoese et al., 2017), *Escherichia coli* (Grad et al., 2012; Holmes et al., 2015; Rusconi et al., 2016), and *Listeria monocytogenes* (Jackson et al., 2016; Kwong et al., 2016; Chen et al., 2017a; Chen et al., 2017b; Moura et al., 2017). However, as demonstrated here and elsewhere (Pightling et al., 2014; Hwang et al., 2015; Pightling et al., 2015; Katz et al., 2017; Sandmann et al., 2017), the choice of variant calling pipeline can influence the identification of SNPs in WGS data. This can be particularly problematic for outbreak and cluster detection in bacterial pathogen surveillance: despite the issues that come with using pairwise SNP difference cutoffs to determine which isolates are included and excluded in an outbreak or cluster (McCloskey and Poon, 2017), SNP thresholds are currently widely used to make initial decisions on the inclusion or exclusion of isolates in a given outbreak (Taylor et al., 2015; Gymoese et al., 2017; Mair-Jenkins et al., 2017; Walker et al., 2018). In such scenarios, just a few SNPs can be the deciding factor in whether a bacterial pathogen is included or excluded as part of an outbreak or cluster (Katz et al., 2017), rendering the choice of variant calling method as non-trivial. Choosing an appropriate variant calling pipeline can be particularly challenging for pathogens where there are limited data and expertise with WGS (e.g., as is currently the case with *B. cereus*).

As demonstrated here, the choice of variant calling pipeline can greatly influence the number of core SNPs identified in *B. cereus* group isolates associated with a foodborne outbreak. In the case of a multi-clade outbreak, this effect can be magnified: naively calling variants in isolates that span multiple *B. cereus* group clades in aggregate can lead to orders of magnitudes of difference in the number of core SNPs identified by different variant calling pipelines/reference genome combinations. In a multi-clade outbreak scenario, it is essential to note that one is effectively dealing with genomic data from *multiple species* (i.e., ANI < 95), making it impossible to find a reference genome that is closely related to all isolates in the outbreak. In the case of some reference-based pipelines that are specifically tailored to identify variants in bacterial isolates from outbreaks (e.g., CFSAN, which is not suited for bacteria differing by more than a few hundred SNPs), calling variants in multiple clades or within a distant reference genome is inappropriate (Davis et al., 2015). Thus, querying outbreak isolates from multiple clades in aggregate using reference-based variant calling methods should be avoided. Furthermore, the results presented here showcase the value of employing single- and/or multi-locus typing approaches prior to variant calling, either via Sanger sequencing or *in silico* using tools such as BTyper, as they can aid the design of downstream bioinformatics analyses (e.g., reference genome selection, data partitioning by clade).

When the three clade IV isolates were excluded from analyses, leaving only the emetic clade III isolates, the selection of reference genome caused fewer core SNP discrepancies than choice of variant calling pipeline, provided the reference genome was “similar” to the genomes analyzed. While the selection of a reference genome for reference-based variant calling is not trivial (Pightling et al., 2014; Olson et al., 2015), reference-based variant calling using a closed chromosome (*B. cereus* str. AH187) and a draft genome (FOOD_10_19_16_RSNT1_2H_R9-6393) from two isolates that were closely related to or among the emetic clade III isolates sequenced in this outbreak produced nearly identical results in terms of the number and identity of SNPs detected. Both reference genomes were identical to the emetic clade III outbreak isolates sequenced here in terms of *panC* clade, *rpoB* AT, MLST ST, and virulotype. Additionally, the closed chromosome and draft genome had ANI values of > 99.8 and 99.9 relative to all emetic clade III outbreak isolates, respectively. Similar findings have been observed in *Salmonella enterica* serovar Heidelberg (Usongo et al., 2018), suggesting that either closed genomes or high-quality draft genomes are adequate for reference-based SNP calling, provided both are similar enough to the outbreak strains being queried.

With regard to differences in the number of core SNPs identified in the 30 emetic clade III isolates using different variant calling pipelines, the pipelines that used assembled genomes as input (kSNP3 and Parsnp) produced higher numbers of core SNPs than their counterparts that relied on short Illumina reads. Additionally, both kSNP3 and Parsnp produced core SNPs that produced topologically similar phylogenies. kSNP3 employs a reference-free *k-*mer based SNP calling approach (Gardner and Hall, 2013; Gardner et al., 2015), while Parsnp uses a reference-based core genome alignment approach (Treangen et al., 2014), and both are useful for calling variants in large data sets. These approaches are also valuable when reads are not available for SNP calling (Olson et al., 2015), as demonstrated here by the comparison of outbreak genomes with publicly-available genomes: core SNPs obtained using both kSNP3 and Parsnp were able to consistently produce phylogenies in which the 30 emetic isolates from this outbreak formed a well-supported clade among all emetic ST 26 *B. cereus* group genomes. However, kSNP3 has been shown to lack specificity relative to other pipelines (i.e., CFSAN, LYVE-SET) when differentiating outbreak isolates from non-outbreak isolates for *L. monocytogenes, E. coli,* and *S. enterica* (Katz et al., 2017). Here, the CFSAN and LYVE-SET pipelines identified similar SNPs that produced highly congruent phylogenies. This is unsurprising, considering both the CFSAN and LYVE-SET pipelines were designed specifically for identifying SNPs in closely-related strains from outbreaks (Katz et al., 2017), and both employ the most stringent filtering criteria of all pipelines tested here.

### 4.3 As WGS becomes routinely integrated into food safety, clinical, and epidemiological realms, it is likely that the number of illnesses attributed to *B. cereus* will increase

Here, we offer the first description of a foodborne outbreak caused by *B. cereus* group species to be characterized using WGS, and we provide a glimpse into the genomic variation one might expect within an emetic clade III *B. cereus* outbreak using several different variant calling pipelines. However, our ability to query emetic clade III genomes outside of this outbreak is limited by the lack of publicly-available genomic data and metadata from emetic isolates. Of the 2,156 *B. cereus* group genomes available in NCBI’s RefSeq database in March 2018, only 29 were from clade III and possessed the *cesABCD* operon, 25 of which belonged to ST 26. While not ideal, this is an improvement, as there were only 19 emetic clade III genomes available in NCBI’s Genbank database in April 2017 (Carroll et al., 2017). As more *B. cereus* group WGS data—particularly, data from emetic *B. cereus* group isolates—become publicly available, more outbreaks and clusters are likely to be resolved in tandem, a phenomenon that has been observed for *L. monocytogenes* (Jackson et al., 2016). Additionally, variant calling and cluster/outbreak detection methods for characterizing *B. cereus* group isolates from foodborne outbreaks can be further refined and optimized as more data and metadata are available for clinical and non-clinical isolates.

## 5. Author Contributions

LC performed computational analyses; MM, LM, ND, and JC performed microbiological experiments. DN provided and interpreted epidemiological data. MW and JK conceived the study. LC, MW, and JK co-wrote the manuscript.

## 6. Funding

This material is based on work supported by the National Science Foundation Graduate Research Fellowship Program under grant no. DGE-1144153. This work was supported also by the USDA National Institute of Food and Hatch Appropriations under Project #PEN04646 and Accession #1015787.

## 7. Conflict of Interest

The authors declare that the research was conducted in the absence of any commercial or financial relationships that could be construed as a potential conflict of interest.

## 8. Acknowledgments

The authors would like to acknowledge the Wadsworth Center Tissue Culture & Media Core for providing the media used in this work, and Dr. Joshua Lambert from The Pennsylvania State University for providing tissue culture laboratory facility and advising.

